# Enhancing Antibody Language Models with Structural Information

**DOI:** 10.1101/2023.12.12.569610

**Authors:** Justin Barton, Jacob D. Galson, Jinwoo Leem

## Abstract

The central tenet of molecular biology is that a protein’s amino acid sequence determines its three-dimensional structure, and thus its function. However, proteins with similar sequences do not always fold into the same shape, and vice-versa, dissimilar sequences can adopt similar folds. In this work, we explore antibodies, a class of proteins in the immune system, whose local shapes are highly unpredictable, even with small variations in their sequence. Inspired by the CLIP method [1], we propose a multimodal contrastive learning approach, contrastive sequence-structure pre-training (CSSP), which amalgamates the representations of antibody sequences and structures in a mutual latent space. Integrating structural information leads both antibody and protein language models to show better correspondence with structural similarity and improves accuracy and data efficiency in downstream binding prediction tasks. We provide an optimised CSSP-trained model, AntiBERTa2-CSSP, for non-commercial use at https://huggingface.co/alchemab.

## 1 Introduction

A protein’s amino acid sequence contains all the information that is necessary to fold it into its three-dimensional structure [2]. The protein’s structure determines its function, meaning that the protein’s sequence contains almost all the information necessary to understand a protein’s function. One strategy that has proven to be very powerful at learning the relationship between sequence and function has been the development of protein language models (PLMs). PLMs process large datasets of protein sequences (e.g. UniRef50) via self-supervised learning to understand the ties between a protein’s sequence, structure, and function [3–5]. However, one class of proteins has proven to be particularly challenging for PLMs due to their extraordinary sequence diversity: antibodies [5–8].

Antibodies are proteins of the immune system. They have a characteristic Y-shape, comprised of two pairs of two protein chains: a heavy chain and a light chain. The combination of heavy-light chain pairs, along with diversification mechanisms such as V(D)J recombination and somatic hypermutation, allow antibodies to reach a theoretical diversity of over 10^15^ possible variants in humans [9, 10]. Much of this variation is concentrated in local stretches of the amino acid sequence, known as the complementarity determining regions (CDRs); the CDRs comprise most of the antibody binding site. In particular, the third CDR of the heavy chain (CDRH3) has the most important role in binding, and it is also the most polymorphic in terms of sequence and structure [11, 12]. Small changes in the CDRH3 sequence can dramatically alter its shape and thus binding specificity; vice-versa, two antibodies with distant sequences can bind the same region on a target protein [13]. This complexity is challenging to learn from antibody sequences alone, and structural data can play an integral role in understanding antibody function.

As of July 2023, there are only 7,463 antibody Protein Data Bank (PDB) entries in SAbDab; 1,085 of these are high-quality human antibody entries with some redundancy (i.e. multiple antibody structures per PDB entry) [14]. While structural modelling tools can be used to fill in the gaps of antibody sequence space, even the fastest prediction tools cannot match the scale of the antibody repertoire. For instance, a typical human antibody repertoire dataset contains 100,000 sequences per individual [15]. Even if a structure prediction tool required only one second per antibody structure [16], this would require more than one day of compute per human repertoire sample. Given this challenging scale, it would be advantageous to have a foundation model that combines both sequence and structural information to capture the nuances of antibody sequence-structure variation, while achieving the inference scalability of a language model.

In this work, we exploit the Contrastive Language-Image Pre-training (CLIP) architecture to update antibody sequence representations with structural information [1]. Our Contrastive Sequence-Structure Pre-training (CSSP) approach involves updating a sequence encoder, while using a frozen, pre-trained structure encoder. Previous approaches in protein multimodal learning have updated both sequence and structure encoders, often investigating how sequence encoders can enrich structure encoders [17–19]. These protein multimodal models have relied on having both sequence and structure data as input during inference. Given the sparsity of available structures, this would likely necessitate using a predicted protein structure [20]. In contrast, we detach the updated sequence encoder for stand-alone use in fine-tuning and inference. We find that our CSSP approach leads to sequence representations that better correspond with antibody structural similarity, and improve accuracy and data efficiency in downstream prediction tasks.

## 2 Methods

### 2.1 Datasets

A total of 779.4 million human antibody sequences were used for pre-training an antibody-specific sequence encoder model, AntiBERTa2. Further details on data processing are described in Appendix A.1.

For multimodal pre-training, we used 1,554 of the human antibody structures downloaded from SAbDab on July 18th, 2023, split into 1,237 training, 155 validation, and 162 test structures [14]. We also used 1.33M model antibody structures, predicted by ABodyBuilder2, to complement the experimental data [21]. For each prediction, ABodyBuilder2 estimates the prediction error per residue. Additional details of structural data filtering and processing are described in Appendix A.1.

All unimodal and multimodal models in this work were benchmarked on an antibody-antigen binding dataset comprised of trastuzumab variants screened for binding to human epidermal growth factor receptor 2 (HER2) [22]. The sizes and stratification details are described in Appendix A.1.

### 2.2 AntiBERTa2 unimodal pre-training

AntiBERTa2 is based on the RoFormer architecture, a bi-directional transformer encoder model with rotary position embeddings [3, 23]. AntiBERTa2 can accept either unpaired (i.e. heavy chain only or light chain only) or paired antibody sequences, where chains are separated by a [SEP] token. AntiBERTa2 was pre-trained on NVIDIA’s DGX SuperCloud (Cambridge-1) using 48 A100 GPUs via masked language modelling. Hyperparameters for AntiBERTa2 are described in Appendix Table A1.

### 2.3 Contrastive sequence-structure pre-training

Our contrastive pre-training approach follows the CLIP procedure [1]. In brief, CLIP minimises

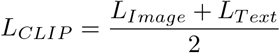

across *N* image-text pairs, where

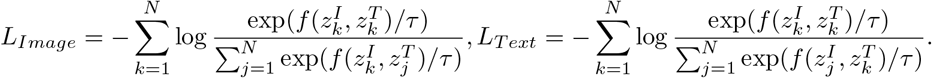

Here, 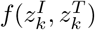 represents the cosine similarity between the k^th^ text embedding, 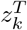, and the k^th^ image embedding 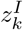, scaled by a learnable temperature parameter *τ*. For CSSP, we analogised sequence embeddings as text embeddings, and structure embeddings as image embeddings.

CSSP was performed with three different PLMs as antibody sequence encoders: AntiBERTa2, ESM2-650M, and AntiBERTy [3, 6]. For each unimodal language model, [CLS] pooling was used to generate the sequence embedding. In all cases, ESM-IF1 was used as the structure encoder [24]. Briefly, ESM-IF1 is a pre-trained geometric vector perceptron model that is linked to a 12-layer transformer encoder-decoder model (6 encoder layers, 6 decoder layers). For our work, we froze the ESM-IF1 model, and average pooled the antibody structure embeddings from the transformer encoder, following the procedure used in the public ESM Google Colab notebook. Hyperparameters for CSSP are described in Appendix Table A2.

### 2.4 Prediction of structural similarity

Structural similarity between two antibody variable regions (Fv) was defined as the root-mean square deviation (RMSD) between backbone atoms (N, C*α*, C, and O). To align antibodies with different sequence lengths, we aligned antibodies based on their IMGT numbering. CDRH3 loop structure similarity was defined as the RMSD between backbone atoms in the IMGT-defined CDRH3 loops. We implemented the dynamic time warp algorithm for length-independent CDRH3 loop alignment, as previously described [25]. Z-score normalization was then applied to the RMSDs.

The latent structural information in pre- and post-CSSP language model embeddings was quantified by predicting pairwise structural similarities using a regression head applied to the [CLS] embeddings of antibody sequences. Specifically, the [CLS] token embeddings of both sequences were concatenated, along with the difference between them, i.e. concat(*z*_*j*_, *z*_*k*_, | *z*_*k*_ − *z*_*j*_ |), in a manner similar to SBERT [26].

### 2.5 Benchmarking on antigen binding

Pre- and post-CSSP PLMs were benchmarked using HER2 binding prediction as a binary classification task. To enable a robust comparison of the resulting sequence embeddings, we froze each model’s sequence encoder weights and only allowed weight updates in the classification head.

## 3 Results

### 3.1 Multimodal training enhances structural awareness of language models

Previous work has shown that PLMs learn aspects of protein structure following pre-training. For example, self-attention scores of transformer models correlate with inter-residue contacts in experimental protein structures [7, 27–29]. However, it is unclear if language model embeddings capture the complex dynamics of structural similarity. Thus, we investigated the relationship between sequence embeddings and antibody structural similarity (Appendix Figure A1).

Before CSSP, we find that embeddings from antibody-specific language models, specifically AntiBERTy and AntiBERTa2, contain more information about antibody structural similarity than the general protein language model ESM2-650M. For all models, applying CSSP increases the structural information content of the embeddings as measured by the correlation of predicted and actual structural similarity. Overall, AntiBERTa2-CSSP embeddings contain the most antibody structure information, with predictions’ Pearson correlation coefficients of 0.706 to Fv similarity, and 0.576 to CDRH3 similarity. One interesting observation is that CSSP was most beneficial for ESM2-650M; this is to be expected as ESM2-650M is not antibody-specific to begin with, and thus CSSP with antibody structures and sequences is effectively making it more of an antibody-specific model.

### 3.2 Contrastive sequence-structure pre-training improves antigen binding prediction

In antibody discovery, the main property, or ‘function’ of interest is an antibody’s binding to its cognate antigen. Thus, we benchmarked AntiBERTa2, ESM2-650M, and AntiBERTy for antigen binding prediction, then compared each model’s performance after CSSP pre-training. We assess models on a dataset of over 20,000 trastuzumab variants binding HER2 [22] (see Appendix A.1).

The CSSP method improves all three models’ predictions of HER2 binding (Figure 1, Appendix Table A3). The mean area under the receiver operating curve (AUROC) score improves each epoch up to 5 epochs of CSSP, after which performance plateaus. Our structural training set contains no relatives of trastuzumab, suggesting that this result is not an artefact of data leakage. Instead, structural information appears to refine the model’s representations for improved binding prediction.

**Figure 1.**
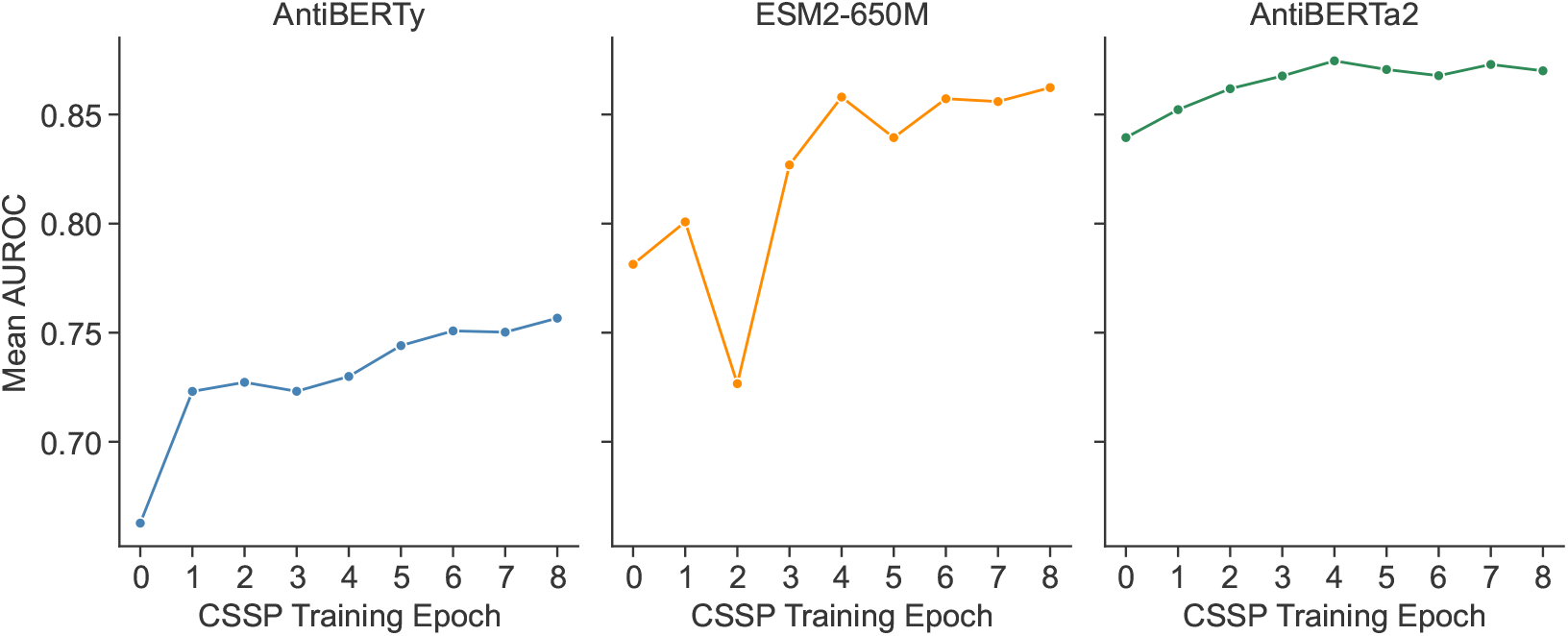
Contrastive sequence-structure pretraining (CSSP) with experimental structural data improves model performance. The mean area under the receiver operating curve (AUROC) score improves per epoch, saturating after 5 epochs. Epoch 0 is the original unimodal model prior to CSSP.

It is often impractical to gather large volumes of antibody-antigen binding data due to cost and time constraints. One of the benefits of multimodal models is their capacity to improve data efficiency in fine-tuning [1, 30]. Thus, we conducted a data ablation study, subsampling the HER2 training set from 0.1%–100%, while using the same test set (Figure 2, Appendix Table A4). For example, at 0.1% of the data, which represents 19 training sequences, AntiBERTa2 achieves an AUROC=0.590; after CSSP, its AUROC improves to 0.686. Once 5% of the training data is used (912 antibodies), AntiBERTa2-CSSP achieves an AUROC of 0.817. For the unimodal AntiBERTa2, this level of performance is only achieved when 40% of the training data becomes available, indicating an 8-fold improvement in efficiency. AntiBERTy and ESM2 both show higher AUROC after CSSP pre-training, with 8-fold and 2-fold efficiency gains respectively. This highlights the flexibility of CSSP approach to any unimodal PLM.

**Figure 2.**
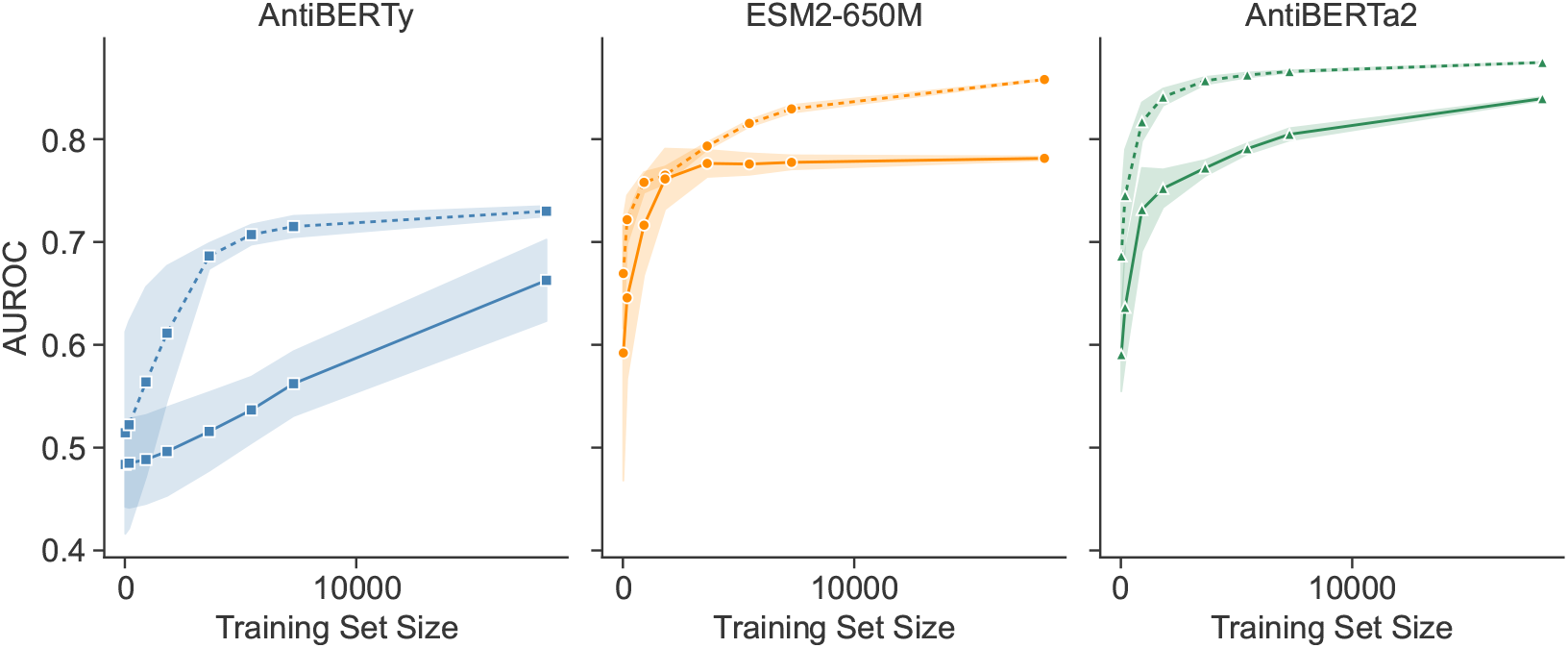
CSSP with experimental structure data improves few-shot binding prediction performance. Solid lines represent the original unimodal model. Dashed lines represent the same models after CSSP pre-training. Shaded areas represent standard deviations using five different random seeds.

Multimodal models consistently outperform their unimodal counterparts across different data ablations, suggesting that structural pre-training fundamentally changes the representations in a way that aids binding prediction. In addition, we note that the standard deviations of AUROC scores across the different data ablations are often reduced after CSSP. These results reflect the value of structural pre-training in low-data settings that are common in antibody discovery. In effect, CSSP provides a means to reduce the burden on wet lab scientists and accelerate antibody engineering campaigns.

### 3.3 Exercise caution when using predicted structures

Due to the relative paucity of publicly available structural data in the PDB, it can be desirable to use predicted structures to increase training datasets [24]. Our work demonstrates that modelled structures do not contribute novel information to CSSP. While better than having no structural information, model structures are not as valuable as crystal structures, even with ten-fold more models (Figure 3A). We also report that combining models with experimental structures provides no additional benefit in CSSP (Figure 3B). Finally, when using half the available experimental structural data, this leads to an expected drop in performance compared to using the full set of 1,237 structures. However, this gap is not recovered by adding predicted structures (Figure 3C). This indicates that it is not simply the volume of structural data, but the information content that has a beneficial impact.

**Figure 3.**
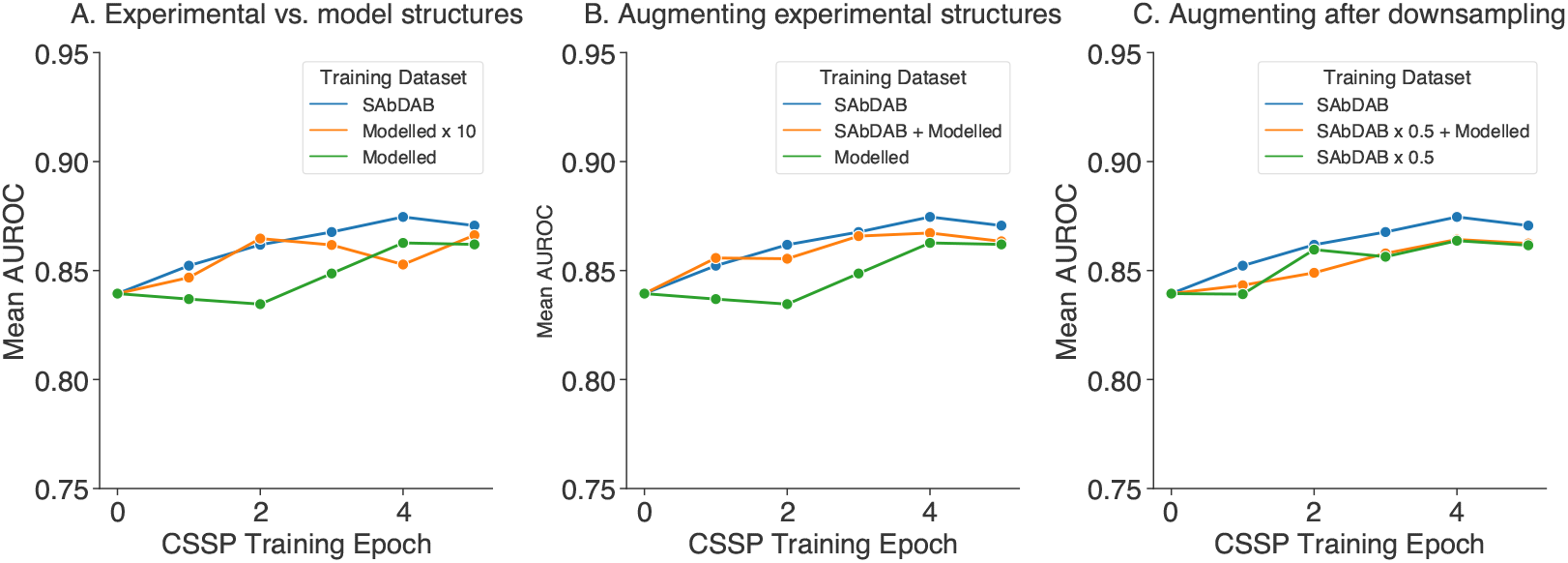
Model structures do not contribute novel information to CSSP. Epoch 0 is AntiBERTa2 before CSSP pre-training. (A) AntiBERTa2-CSSP with 1,237 experimental structures (blue) has higher performance compared to AntiBERTa2-CSSP pre-trained on 1,237 model structures (green) and 12,370 model structures (orange). (B) Combining predicted structures with experimental structures (orange) does not provide synergistic benefit. (C) Halving the experimental training data used in AntiBERTa2-CSSP (green) leads to a performance drop; this is not rescued by models (orange).

## 4 Conclusion

In this study, we demonstrate how structural information can be imbued into PLMs by using contrastive sequence-structre pre-training. We find that CSSP increases the amount of structural information encoded in PLM embeddings, leading to stronger correlations to both global and CDRH3 structure variation. An exciting advantage of CSSP is its impact on few-shot antigen binding prediction. Even with a small dataset of experimental structures, CSSP can improve performance and increase data efficiency, emphasising the importance of structural knowledge in understanding function. A further benefit of CSSP is its flexibility: in principle, CSSP can be used with any PLM. Our multimodal learning approach offers a pragmatic solution to antibody engineering, and it can be especially helpful in resource-limited environments.

## Acknowledgments and Disclosure of Funding

We would like to thank Aretas Gaspariunas for standardising the structural data collection pipeline. We also extend our thanks to James H.R. Farmery and Harry Dobson for maintaining our compute infrastructure. A special thanks goes to our wonderful lab scientists, Jorge Dias, David A. Yadin, Francesca L. Nice, Danielle H. Minns, Chelsea Povall, Sara Valle Tomas, and Olivia Snudden for their sequencing work to support this study. We would also like to acknowledge Konrad Krawczyk of NaturalAntibody for providing the 1.33 million antibody model structures. The authors would finally like to thank Bruno Trentini, Hassan Sirelkhatim, Christian Dallago, Maxine Kennedy, Lee Carter, and the wider NVIDIA team for their support.

## A Appendix

### A.1 Dataset preparation

For pre-training AntiBERTa2, we collected 1.47 billion unpaired (i.e. heavy chain or light chain only) human antibodies from the Observed Antibody Space (OAS) database on 23^rd^ February 2023. We also included 70 million unpaired human antibodies from a proprietary dataset. We used Linclust to cluster antibodies at 90% sequence identity, leading to a dataset of 821.2 million unpaired antibodies.

In parallel, we collected 1.5 million paired human antibody sequences [31], and a further 1.4 million from an in-house experiment. Due to the relative paucity of paired data, we implemented a 99% sequence identity cut-off, leading to a total of 2.5 million paired antibodies. The ensemble of unpaired and paired data were stratified into a 95:5 train:evaluation ratio for masked language modelling: 779.4 million sequences were in the training set, and 44.3 million in the evaluation set.

Antibody structures were downloaded from SAbDab on the 18^th^ July 2023. Structures resolved by either X-ray crystallography or cryo-electron microscopy were filtered for resolution of 2.5Å or better; structures with missing coordinates in the CDR residues were removed. We also removed antibodies that were single-chain variable fragments and omitted non-human antibodies. Since antibodies with identical amino acid sequences can have slight variations in their structure, we retained a redundant dataset of structures, leading to a final set of 1,554 antibody structures from 995 PDB entries. Structures were stratified by their PDB code in an ∼ 80:10:10 ratio to train, evaluate, and test CSSP. In total, we used 1,237 structures in the training, 155 in evaluation, and 162 in the test set. Modelled structures were predicted using ABodyBuilder2 for paired B cell receptor sequences [21, 31], and we used a random subset of model structures for our analyses.

For benchmarking, a dataset comprising 39,108 variants of trastuzumab that were screened for binding to the human epidermal growth factor receptor 2 (HER2) protein were downloaded [22]. Briefly, the experiment used hybridomas expressing either complementarity determining region (CDR) H3 or CDRL3 variants, and antigen binding was confirmed via cell sorting. Antibodies that do not bind the antigen, i.e. negatives, were randomly under-sampled; the remaining antibodies were randomly split into disjoint train, test, and validation sets as done previously. In total, 18,223, 2,278, and 2,278 sequences, respectively.

**Figure A1:**
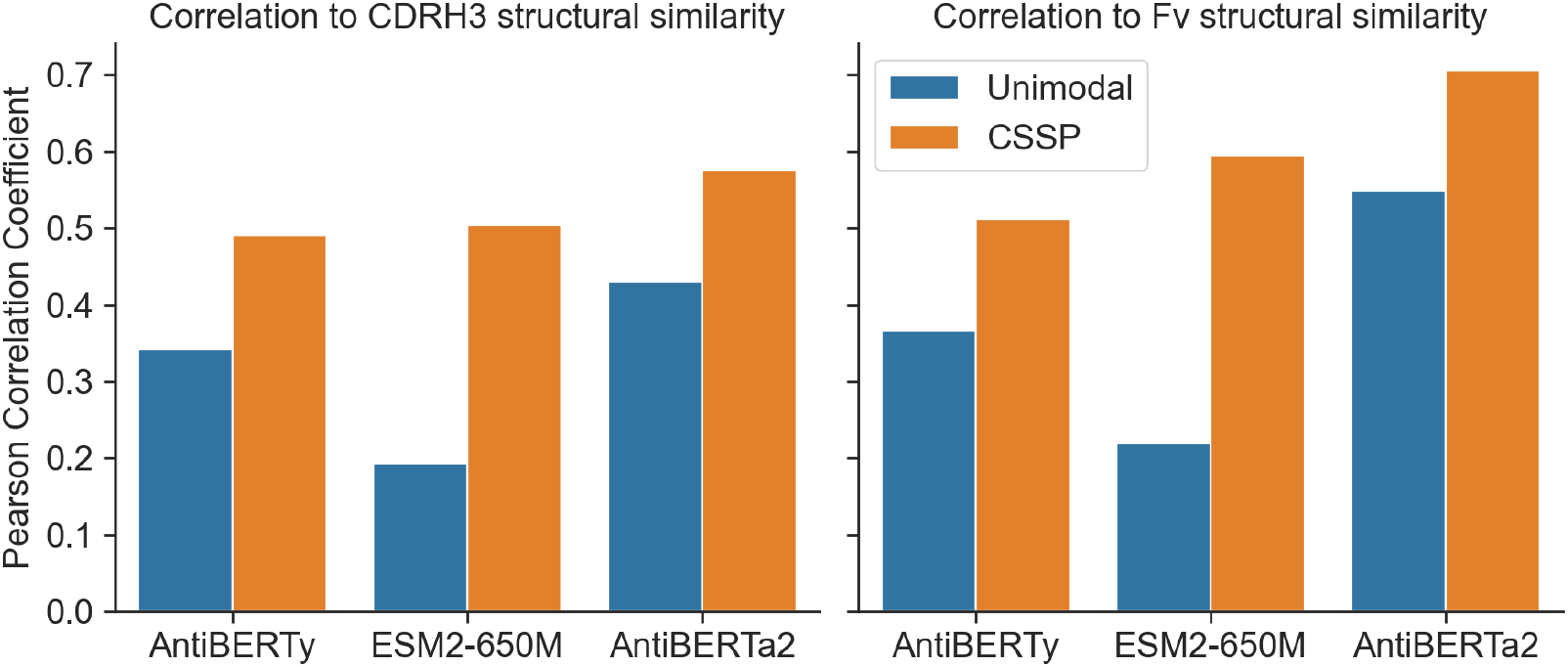
Pearson correlation coefficient between predicted and actual structural similarity before and after CSSP. Structural similarity is predicted from embeddings using a regression head as described in the methods. The calculation of structural similarity for the CDRH3 loop and the entire antibody variable domain (Fv) is also described in the methods. CSSP increases the structural information content of all unimodal language models’ embeddings.

**Table A1:** Hyperparameters for pre-training AntiBERTa2.

|  |  |
| --- | --- |
| **Number of layers** | 16 |
| **Number of attention heads** | 16 |
| **Embedding dimension** | 1024 |
| **Feed-forward dimension** | 4096 |
| **Peak learning rate** | 8e-5 |
| **Learning rate schedule** | Linear |
| **Number of steps** | 260k (40k warmup) |
| **Optimizer** | AdamW |
| **Weight decay** | 0.01 |

**Table A2:**
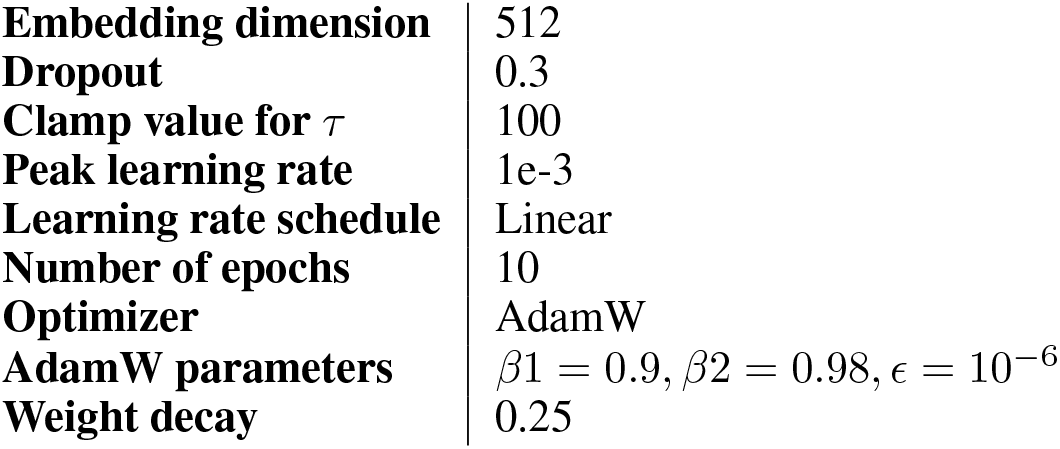
Hyperparameters for CSSP.

**Table A3:** HER2 Binding Prediction Performance.

| **Model** | **AUROC** |  | **AUPR** |  | **F1** |  |
| --- | --- | --- | --- | --- | --- | --- |
|  | *mean* | *std* | *mean* | *std* | *mean* | *std* |
| AntiBERTy | 0.663 | 0.040 | 0.641 | 0.039 | 0.669 | 0.002 |
| AntiBERTy-CSSP | 0.730 | 0.005 | 0.707 | 0.005 | 0.700 | 0.006 |
| ESM2-650M | 0.781 | 0.002 | 0.739 | 0.002 | 0.706 | 0.031 |
| ESM2-650M-CSSP | 0.858 | 0.001 | 0.853 | 0.001 | 0.781 | 0.001 |
| AntiBERTa2 | 0.839 | 0.002 | 0.827 | 0.001 | 0.754 | 0.006 |
| AntiBERTa2-CSSP | 0.875 | 0.001 | 0.869 | 0.001 | 0.807 | 0.001 |

**Table A4:** HER2 Binding Prediction Performance (AUROC)

| Model | 0.1% $n=19$ | 1% $n=183$ | 5% $n=912$ | 10% $n=1823$ | 20% $n=3645$ | 30% $n=5467$ | 40% $n=7290$ | 100% $n=18223$ |
| --- | --- | --- | --- | --- | --- | --- | --- | --- |
| AntiBERTy | $0.484 \pm 0.041$ | $0.485 \pm 0.043$ | $0.488 \pm 0.043$ | $0.496 \pm 0.043$ | $0.516 \pm 0.038$ | $0.537 \pm 0.032$ | $0.562 \pm 0.031$ | $0.663 \pm 0.040$ |
| AntiBERTy-CSSP | $0.514 \pm 0.098$ | $0.522 \pm 0.101$ | $0.564 \pm 0.092$ | $0.611 \pm 0.066$ | $0.686 \pm 0.013$ | $0.707 \pm 0.010$ | $0.715 \pm 0.010$ | $0.730 \pm 0.005$ |
| ESM2-650M | $0.592 \pm 0.124$ | $0.646 \pm 0.079$ | $0.716 \pm 0.049$ | $0.761 \pm 0.029$ | $0.776 \pm 0.013$ | $0.776 \pm 0.010$ | $0.777 \pm 0.007$ | $0.781 \pm 0.002$ |
| ESM2-650M-CSSP | $0.669 \pm 0.054$ | $0.722 \pm 0.024$ | $0.758 \pm 0.010$ | $0.765 \pm 0.008$ | $0.793 \pm 0.004$ | $0.815 \pm 0.003$ | $0.829 \pm 0.004$ | $0.858 \pm 0.001$ |
| AntiBERTa2 | $0.590 \pm 0.036$ | $0.636 \pm 0.053$ | $0.731 \pm 0.040$ | $0.752 \pm 0.018$ | $0.772 \pm 0.008$ | $0.791 \pm 0.005$ | $0.805 \pm 0.006$ | $0.839 \pm 0.002$ |
| AntiBERTa2-CSSP | $0.686 \pm 0.054$ | $0.745 \pm 0.045$ | $0.817 \pm 0.019$ | $0.841 \pm 0.008$ | $0.857 \pm 0.004$ | $0.862 \pm 0.002$ | $0.866 \pm 0.001$ | $0.875 \pm 0.001$ |

## References

[1] A. Radford, J. W. Kim, C. Hallacy, A. Ramesh, G. Goh, S. Agarwal, G. Sastry, A. Askell, P. Mishkin, J. Clark, G. Krueger, and I. Sutskever, “Learning Transferable Visual Models From Natural Language Supervision,” arXiv, 2 2021.

[2] B. Kuhlman and P. Bradley, “Advances in protein structure prediction and design,” Nature Reviews Molecular Cell Biology, vol. 20, no. 11, pp. 681–697, 2019.

[3] Z. Lin, H. Akin, R. Rao, B. Hie, Z. Zhu, W. Lu, N. Smetanin, R. Verkuil, O. Kabeli, Y. Shmueli, A. d. S. Costa, M. Fazel-Zarandi, T. Sercu, S. Candido, and A. Rives, “Evolutionary-scale prediction of atomic-level protein structure with a language model,” Science, vol. 379, no. 6637, pp. 1123–1130, 2023.

[4] A. Elnaggar, M. Heinzinger, C. Dallago, G. Rehawi, Y. Wang, L. Jones, T. Gibbs, T. Feher, C. Angerer, M. Steinegger, D. Bhowmik, and B. Rost, “ProtTrans: Toward Understanding the Language of Life Through Self-Supervised Learning,” IEEE Transactions on Pattern Analysis and Machine Intelligence, vol. 44, no. 10, pp. 7112–7127, 2022.

[5] E. Nijkamp, J. Ruffolo, E. N. Weinstein, N. Naik, and A. Madani, “ProGen2: Exploring the Boundaries of Protein Language Models,” arXiv, 2022.

[6] J. A. Ruffolo, J. J. Gray, and J. Sulam, “Deciphering antibody affinity maturation with language models and weakly supervised learning,” arXiv, 2021.

[7] J. Leem, L. S. Mitchell, J. H. Farmery, J. Barton, and J. D. Galson, “Deciphering the language of antibodies using self-supervised learning,” Patterns, vol. 3, no. 7, p. 100513, 2022.

[8] B. Chen, X. Cheng, Y.-a. Geng, S. Li, X. Zeng, B. Wang, J. Gong, C. Liu, A. Zeng, Y. Dong, J. Tang, and L. Song, “xTrimoPGLM: Unified 100B-Scale Pre-trained Transformer for Deciphering the Language of Protein,” bioRxiv, 2023.

[9] G. Georgiou, G. C. Ippolito, J. Beausang, C. E. Busse, H. Wardemann, and S. R. Quake, “The promise and challenge of high-throughput sequencing of the antibody repertoire,” Nature Biotechnology, vol. 32, no. 2, pp. 158–168, 2014.

[10] A. R. Rees, “Understanding the human antibody repertoire,” mAbs, vol. 12, no. 1, p. 1729683, 2020.

[11] C. Regep, G. Georges, J. Shi, B. Popovic, and C. M. Deane, “The H3 loop of antibodies shows unique structural characteristics,” Proteins: Structure, Function, and Bioinformatics, vol. 85, no. 7, pp. 1311–1318, 2017.

[12] M. L. Fernández-Quintero, J. R. Loeffler, J. Kraml, U. Kahler, A. S. Kamenik, and K. R. Liedl, “Characterizing the Diversity of the CDR-H3 Loop Conformational Ensembles in Relationship to Antibody Binding Properties,” Frontiers in Immunology, vol. 9, p. 3065, 2019.

[13] A. Kovaltsuk, K. Krawczyk, J. D. Galson, D. F. Kelly, C. M. Deane, and J. Trück, “How B-Cell Receptor Repertoire Sequencing Can Be Enriched with Structural Antibody Data,” Frontiers in Immunology, vol. 8, p. 1753, 2017.

[14] J. Dunbar, K. Krawczyk, J. Leem, T. Baker, A. Fuchs, G. Georges, J. Shi, and C. M. Deane, “SAbDab: the structural antibody database,” Nucleic Acids Research, vol. 42, no. D1, pp. D1140–D1146, 2014.

[15] J. D. Galson, S. Schaetzle, R. Bashford-Rogers, M. Raybould, A. Kovaltsuk, G. Kilpatrick, R. Minter, D. Finch, J. Dias, L. James, G. Thomas, W. Lee, J. Betley, O. Cavlan, A. Leech, C. Deane, J. Seoane, C. Caldas, D. Pennington, P. Pfeffer, and J. Osbourn, “Deep Sequencing of B Cell Receptor Repertoires From COVID-19 Patients Reveals Strong Convergent Immune Signatures,” Front Immunol, vol. 11, p. 605170, 2020.

[16] J. H. Lee, P. Yadollahpour, A. Watkins, N. C. Frey, A. Leaver-Fay, S. Ra, K. Cho, V. Gligorijević, A. Regev, and R. Bonneau, “EquiFold: Protein Structure Prediction with a Novel Coarse-Grained Structure Representation,” bioRxiv, 2023.

[17] M. Heinzinger, M. Littmann, I. Sillitoe, N. Bordin, C. Orengo, and B. Rost, “Contrastive learning on protein embeddings enlightens midnight zone,” NAR Genomics and Bioinformatics, vol. 4, no. 2, p. lqac043, 2022.

[18] J. Luo and Y. Luo, “Contrastive learning of protein representations with graph neural networks for structural and functional annotations,” Biocomputing, pp. 109–120, 1 2023.

[19] D. Wang, U. L. Abbas, Q. Shao, J. Chen, and D. Xu, “S-PLM: Structure-aware Protein Language Model via Contrastive Learning between Sequence and Structure,” bioRxiv, 2023.

[20] K. K. Yang, H. Yeh, and N. Zanichelli, “Masked Inverse Folding with Sequence Transfer for Protein Representation Learning,” bioRxiv, 2023.

[21] B. Abanades, W. K. Wong, F. Boyles, G. Georges, A. Bujotzek, and C. M. Deane, “ImmuneBuilder: Deep-Learning models for predicting the structures of immune proteins,” Communications Biology, vol. 6, no. 1, p. 575, 2023.

[22] D. M. Mason, S. Friedensohn, C. R. Weber, C. Jordi, B. Wagner, S. M. Meng, R. A. Ehling, L. Bonati, J. Dahinden, P. Gainza, B. E. Correia, and S. T. Reddy, “Optimization of therapeutic antibodies by predicting antigen specificity from antibody sequence via deep learning,” Nature Biomedical Engineering, vol. 5, no. 6, pp. 600–612, 2021.

[23] J. Su, Y. Lu, S. Pan, A. Murtadha, B. Wen, and Y. Liu, “RoFormer: Enhanced Transformer with Rotary Position Embedding,” arXiv, 8 2022.

[24] C. Hsu, R. Verkuil, J. Liu, Z. Lin, B. Hie, T. Sercu, A. Lerer, and A. Rives, “Learning inverse folding from millions of predicted structures,” bioRxiv, 2022.

[25] J. Nowak, T. Baker, G. Georges, S. Kelm, S. Klostermann, J. Shi, S. Sridharan, and C. M. Deane, “Length-independent structural similarities enrich the antibody CDR canonical class model,” mAbs, vol. 8, no. 4, pp. 751–760, 2016.

[26] N. Reimers and I. Gurevych, “Sentence-BERT: Sentence Embeddings using Siamese BERT-Networks,” CoRR, vol. abs/1908.10084, 2019.

[27] J. Vig, A. Madani, L. R. Varshney, C. Xiong, R. Socher, and N. F. Rajani, “BERTology Meets Biology: Interpreting Attention in Protein Language Models,” arXiv, 2020.

[28] D. Prihoda, J. Maamary, A. Waight, V. Juan, L. Fayadat-Dilman, D. Svozil, and D. A. Bitton, “BioPhi: A platform for antibody design, humanization, and humanness evaluation based on natural antibody repertoires and deep learning,” mAbs, vol. 14, no. 1, p. 2020203, 2022.

[29] S. M. Burbach and B. Briney, “Improving antibody language models with native pairing,” arXiv, 2023.

[30] C. Q. Nguyen, D. Pertusi, and K. M. Branson, “Molecule-Morphology Contrastive Pretraining for Transferable Molecular Representation,” arXiv, 6 2023.

[31] D. B. Jaffe, P. Shahi, B. A. Adams, A. M. Chrisman, P. M. Finnegan, N. Raman, A. E. Royall, F. Tsai, T. Vollbrecht, D. S. Reyes, N. L. Hepler, and W. J. McDonnell, “Functional antibodies exhibit light chain coherence,” Nature, vol. 611, no. 7935, pp. 352–357, 2022.

